# Rhizaria in the oligotrophic ocean exhibit clear temporal and vertical variability

**DOI:** 10.1101/2024.05.10.593577

**Authors:** Alex Barth, Leocadio Blanco-Bercial, Rod Johnson, Joshua Stone

## Abstract

Recently studies have shown that Rhizaria, a super-group of marine protists, have a large role in pelagic ecosystems. They are unique in that they construct mineral tests out of silica, calcium carbonate, or strontium sulfate. As a consequence, Rhizaria can have large impacts on the ocean’s cycling of carbon and other elements. However, less is known about Rhizaria ecology or their role in the pelagic food-web. Some taxa, like certain Radiolaria, are mixotrophic, hosting algal symbionts. While other taxa are flux-feeders or even predatory carnivores. Some prior research has suggested that Rhizaria will partition vertically in the water column, likely due to different trophic strategies. However, very few studies have investigated their populations over extended periods of time. In this study, we present data investigating Rhizaria abundance and vertical distribution from over a year of monthly cruises in the Sargasso Sea. This study represents the first quantification of Rhizaria throughout the mesopelagic zone in an oligotrophic system for an extended period of time. We use this data to investigate the hypothesis that Rhizaria taxonomic groups will partition due to trophic mode. We also investigate how their abundance varies in accordance with environmental parameters. Rhizaria abundance was quantified using an Underwater Vision Profiler (UVP5), an in-situ imaging device. Ultimately, we show that different Rhizaria taxa will have unique vertical distribution patterns. Models relating their abundance to environmental parameters have mixed results, yet particle concentration is a common predictive variable, supporting the importance of heterotrophy amongst many taxa.

## Introduction

Rhizaria are an extremely diverse super-group of single-celled organisms consisting of several phyla including Retaria (foraminifera and radiolaria) and Cercozoa. These organisms exist in a wide range of habitats and are widely represented in plankton communities throughout the global ocean. While the taxonomy of these organisms has recently undergone several reclassifications (Biard, 2022a), their presence in ocean ecosystems has been long known to oceanographers. Some of the earliest records of their existence are from oceanographic expeditions in the 19th century (Haeckel, 1887). Rhizaria are unique members of the plankton and protist community because they can reach large sizes (up to several mm in diameter) and they construct intricate mineral skeletons out of either silica, strontium sulfate, or calcium carbonate (Biard, 2022a; Kimoto, 2015; Nakamura and Suzuki, 2015; Suzuki and Not, 2015). Despite their noticeable morphology and global distribution, Rhizaria were largely understudied throughout the 20th century. The bulk of modern plankton research has focused on hard-bodied crustacea which are numerically dominant and easily sampled with nets and preservatives. Fragile organisms like Rhizaria were difficult to adequately study as they can be destroyed through standard zooplankton sampling techniques and preserve poorly. A number of studies in the late 1900s did employ alternative techniques to quantify Rhizaria including diaphragm pumps (Michaels, 1988) or blue-water SCUBA collections (Bijma et al., 1990; Caron et al., 1995; Caron and Be, 1984). However, the bulk of Rhizaria research was constrained to sediment traps or paleontological studies of sediment (Boltovskoy et al., 1993; Takahashi et al., 1983). Only recently has the advent of molecular techniques and in-situ imaging tools ignited a renewed focus on Rhizaria in pelagic ecosystems (Caron, 2016).

The wave of new data on Rhizaria has facilitated an improved understanding of the significance in ocean ecosystem functions. Firstly, taxonomists have been able to greatly refine the understanding of evolutionary relationships amongst these diverse protists (Aurahs et al., 2009; Biard et al., 2015; Cavalier-Smith et al., 2018; Decelle et al., 2013, 2012; rev by Biard, 2022a). DNA metabarcoding studies have revealed insights into the distributional patterns (Biard et al., 2017; Blanco-Bercial et al., 2022; Decelle et al., 2013; Llopis Monferrer et al., 2022; Mars Brisbin et al., 2020; Sogawa et al., 2022), ecological relationships (Decelle et al., 2012; Nakamura et al., 2023), and contribution to biogeochemical fluxes (Guidi et al., 2016; Gutierrez-Rodriguez et al., 2019). Transcriptomic and proteomic approaches also have been used to quantify Rhizaria contribution to community metabolism (Cohen et al., 2023). Yet, despite the excellent taxonomic resolution provided by molecular approaches, they do not provide a truly quantitative metric for estimating Rhizaria abundance or biomass. In-situ imaging tools however, offer the ability to observe organisms in the natural state and quantify their abundance (Barth and Stone, 2024; Ohman, 2019). Biard et al. (2016) utilized in-situ imaging at a global scale to suggest Rhizaria were substantial contributors to the ocean carbon standing stock. While more recent calculations suggest lower carbon contribution (Laget et al., 2024), Rhizaria nonetheless have substantial influences on biogeochemical cycling. Due to their large sizes, ability to concentrate smaller particles and the unique structure of their mineral skeletons, Rhizaria have the potential to massively influence ocean biogeochemical cycling. A number of studies have made large advances in estimating the contribution of Rhizaria to ocean cycling of carbon (Gutierrez-Rodriguez et al., 2019; Ikenoue et al., 2019; Lampitt et al., 2009; Stukel et al., 2018), silica (Biard et al., 2018; Llopis Monferrer et al., 2021), and strontium (Decelle et al., 2013). Still, Rhizaria ecological roles are not well understood (Biard, 2022a). This is a major challenge as it is critical to understand the ecological role of plankton to fully incorporate them into biological oceanographic models.

The ecological role of Rhizaria in plankton communities is complicated due to the fact different taxa can exhibit every different trophic modes. As zooplankton, rhizaria are predominately heterotrophic (Biard, 2022a), yet their feeding modes can be quite varied. Phaeodaria (family Cercozoa) are largely thought to be flux-feeders, collecting and feeding on sinking particles (Nakamura and Suzuki, 2015; Stukel et al., 2019). Alternatively, Retaria can be either exclusively heterotrophic or mixotrophic, utilizing photosynthetic algal symbionts (Anderson, 2014; Decelle et al., 2015). Mixotrophic foraminifera host a variety of endosymbiont partners (Decelle et al., 2015; Lee, 2006), which are thought to support early and adult life stages and significantly contribute to total primary productivity (Kimoto, 2015). Still, foraminifera are omnivorous, possibly even predominately carnivorous, with several studies suggesting that they can be effective predators (Anderson and Bé, 1976; Gaskell et al., 2019), mainly consuming live copepods (Caron and Be, 1984). Radiolaria have several lineages, all of which have some taxa well known to host symbionts (Biard, 2022b). Amongst Radiolaria, arguably the most widespread are Collodaria who can be either large solitary cells or form massive colonies, up to several meters in length (Swanberg and Anderson, 1981). All known Collodaria species host dinoflagellate symbionts (Biard, 2022b) and can contribute substantially to primary productivity, particularly in oligotrophic ocean regions (Caron et al., 1995; Dennett et al., 2002). This Collodaria-symbiont association has been suggested as a reason for their high abundances throughout the photic zone of oligotrophic environments globally (Biard et al., 2017, 2016). A few Acantharea (Radiolaria order) clades host algal symbionts (Biard, 2022b; Decelle et al., 2012), notably with two clades forming an exclusive relationship with *Phaeocystis*. However, globally, Acantharea are less abundant than Collodaria (Biard, 2022a) and contribute less to total primary productivity (Michaels et al., 1995). This may be due to the fact several clades of Acantharea are cyst-forming and strictly heterotrophs (Biard, 2022b; Decelle et al., 2013). Furthermore, Mars Brisbin et al. (2020) documented apparent predation behavior in Acantharea near the surface, suggesting that there may be a larger reliance on carnivory.

Given the high abundances, yet diverse trophic strategies found among Rhizaria taxa, it is reasonable to expect some form of niche partitioning. A number of studies do suggest evidence for vertical zonation between Rhizaria groups according to various trophic strategies. Taxa-specific studies of Radiolaria suggest they may be restricted to the euphotic zone (Boltovskoy, 2017; Michaels, 1988). Although some studies report Acantharea in deeper waters (Decelle et al., 2013; Gutiérrez-Rodríguez et al., 2022), Phaeodaria, alternatively, are generally found in the mesopelagic where photosynthesis cannot occur, but they can feed on sinking particles (Stukel et al., 2018). In an imaging-based study of the whole Rhizaria community, Biard and Ohman (2020) noted clear patterns in vertical zonation which largely corresponded to different trophic roles. In the oligotrophic ocean, Blanco-Bercial et al. (2022) also noted that the protist community, including Rhizaria, partition along an autotroph and mixotroph to heterotroph gradient with increasing depth in the water column. Yet, few studies have made direct attempts to relate rhizaria abundances to environmental factors (Biard and Ohman, 2020). In part, this is due to the fact few studies have been able to sample Rhizaria in the same location over a consistent timeframe (Boltovskoy et al., 1993; Gutiérrez-Rodríguez et al., 2022; Hull et al., 2011; Michaels et al., 1995; Michaels, 1988). Furthermore, no studies have utilized imaging, arguably the best method for quantifying rhizaria, consistently throughout the full mesopelagic. Given this lack of information, there are many unknowns with respect to Rhizaria ecology, seasonality and phenology across different groups.

In this study, we present a comprehensive assessment of large Rhizaria measured for over a year from regularly occurring cruises at monthly intervals. We utilized an in-situ imaging approach to facilitate abundance calculations. With this dataset, we address two critical aims. 1) Quantification of large Rhizaria throughout the epipelagic (0-200m) and mesopelagic (200-1000m) over the course of an annual cycle. These data were collected in the Sargasso Sea, and represents the first study of its kind in an oligotrophic system; and 2) We aim to test the hypothesis that Rhizaria exhibit niche partitioning according to trophic roles. This hypothesis makes several predictions, including vertical zonation, as seen in prior studies, but also that environmental variables related to trophic strategy will explain abundance patterns. Specifically, autotrophic/mixotrophic taxa will correspond to variables related to autotrophy (chl-a concentration, primary productivity, local DO maxima) and other rhizaria will correspond to factors which promote heterotrophy (particle concentration, flux, and local DO minima).

## Methods

### Oceanographic Sampling

Data were collected in collaboration with the Bermuda Atlantic Time-series Study (Lomas et al., 2013; Michaels and Knap, 1996) on board the R/V Atlantic Explorer. Cruises were conducted at approximately monthly intervals. Rhizaria individuals were sampled using the Underwater Vision Profiler 5 [UVP5; Picheral et al. (2010)], a tool which is well established to accurately quantify large Rhizaria (Barth and Stone, 2022; Biard et al., 2016; Biard and Ohman, 2020; Drago et al., 2022; Llopis Monferrer et al., 2022; Panaïotis et al., 2023; Stukel et al., 2019; Stukel et al., 2018). The UVP5 was mounted to the sampling rosette and collected data autonomously on routine casts, from which only the downcast data are utilized. The UVP5 was deployed from June-September 2019 then from October 2020 - January 2022, during which time the BATS region was sampled for 3-5 days at monthly intervals. Casts were filtered to only include data collected in the BATS region, far offshore of Bermuda in the Sargasso Sea (approximately 31.0𝑜𝑁-32.5𝑜𝑁, 64.25𝑜𝑊 -63𝑜𝑊 ; Supplemental Figure 1). In general, casts extended to either 200m, 500m, or 1200m deep, with a few extended into the bathypelagic (4500m). However, Rhizaria were only typically found in large abundances throughout the epipelagic and mesopelagic zones. As such, we limit this study to results from the upper 1000m of the water column.

A variety of biotic and abiotic data were collected during each BATS cruise. Briefly, we will explain the data utilized in this study. The UVP5 provided particle count data at a high-frequency from each cast. Particle concentration was calculated from this data for all particles 184𝜇𝑚 - 450𝜇𝑚. The lower size range was set by what could be reliably sampled by the UVP5’s pixel resolution (>2px; 0.092mm per pixel) and the upper size range is representative of a potential prey field for mesozooplankton (Whitmore and Ohman, 2021). For each UVP cast supporting continuous profiles of the CTD parameters salinity, temperature, and auxiliary CTD channels; Dissolved Oxygen (DO), in-situ chlorophyll fluorescence were measured at 24Hz using the BATS CTD package. On select casts, Niskin bottles were used to collect bacterial abundance estimates (via epifluorescence microscopy) as well as measure inorganic nutrients (𝑁𝑂_3_, and 𝑆𝑖 as silicate/sicilic acid) at discrete depths. On each cruise, flux estimates of total mass, organic carbon, and nitrogen were also collected using sediment traps; in the present study we utilized flux to the mesopelagic as the flux at 200m. Also primary productivity was estimated through measuring 14𝐶 uptake rates from in-situ incubations. For full descriptions of the BATS sampling program and methods, see Knap et al. (1997) and Lomas et al. (2013) for a review. Additionally, data can be viewed online (https://bats.bios.asu.edu/bats-data/).

Environmental data were processed in a variety of ways to match the format of the Rhizaria abundance estimates (see below). CTD data were collected at higher frequency than the UVP (24Hz vs 15Hz respectively), so these data were averaged within matching bins to the UVP5 data. Data from Niskin bottles were first linearly interpolated in depth at 1m resolution then time averaged over the cruise, then subsequently averaged into matching UVP5-sized bins. Primary productivity estimates were totaled within the euphotic zone to represent a “total euphotic productivity”.

### Rhizaria imaging processing and quantification

Individual vignettes of Rhizaria images were identified using the classification platform Ecotaxa (Picheral et al., 2017). Data were pre-sorted utilizing a random-forest classifier and pre-trained learning set. Taxonomic classification were done based on morphology exclusively. While there are sparse taxonomic guides for in-situ images of rhizaria, identification largely relied on descriptions in (Nakamura and Suzuki, 2015; Suzuki and Not, 2015; and Biard and Ohman, 2020). Using the aforementioned sources and publicly available ecotaxa projects, we constructed a guide accessible at: https://thealexbarth.github.io/media//Project_Items/Oligotrophic_Community/ecotaxa_UVP-guide-stone-lab.pdf. Broadly, Rhizaria were classified as Foraminifera, Radiolaria (Acantharea or Collodaria), or as a variety of Phaeodaria families (Figure 1). When identification could not be confidently made between a few candidate taxa, a less specific label was used. As a result, we have data from “unidentified Rhizaria”, which typically were vignettes not distinguishable between Aulacanthidae or Acantharea or “unidentified Phaeodaria”, which are clearly Phaeodaria but not distinguishable into a family.

**Figure 1:**
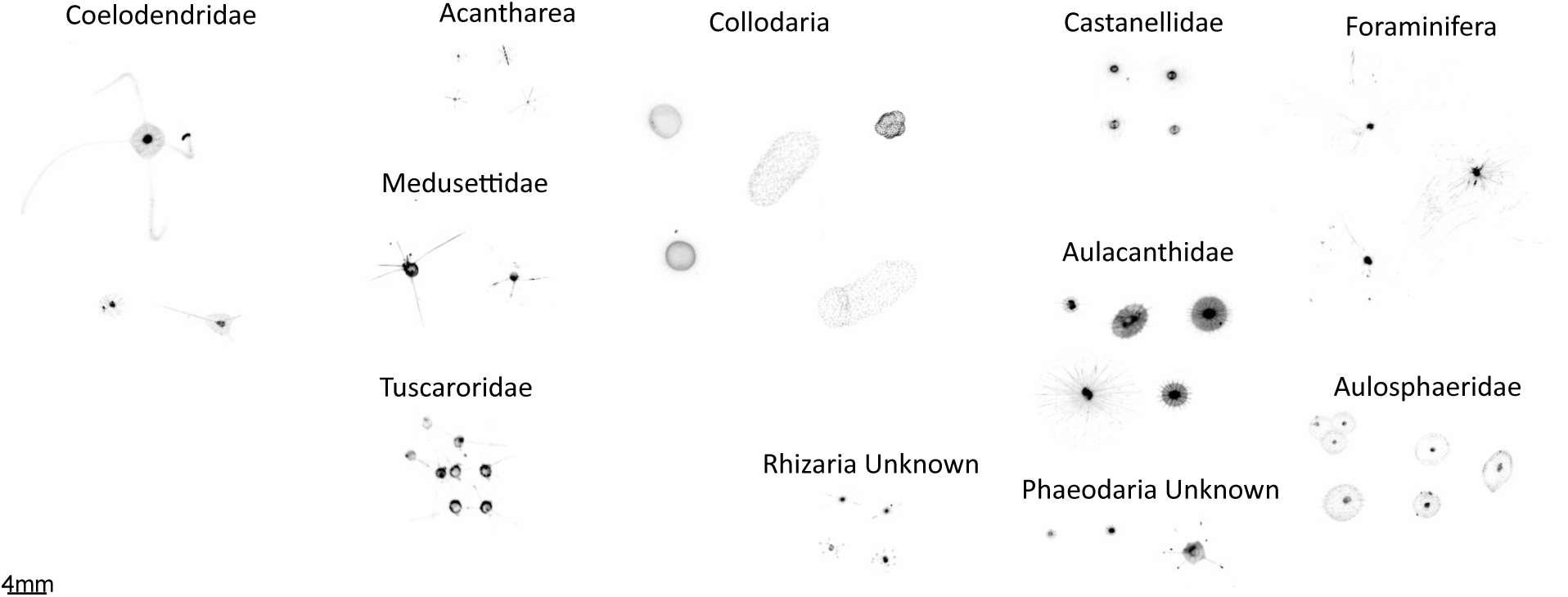
Example images of different Rhizaria taxa. 4mm scale bar shown in lower right. All vignettes are same scale

The UVP5 samples at ∼15Hz rate as it descends the water column and records the exact position of each particle larger than 600𝜇𝑚. However, identified rhizaria ranged from a 934𝜇𝑚 Aulacanthidae cell to a Collodaria colony over 10mm in diameter. To confirm that the UVP5 was sampling adequately across all size ranges, an normalized biomass size spectrum (NBSS) slope was constructed to identify a drop-off which would indicate poor-sampling at the small size range (Barth and Stone, 2024; Lombard et al., 2019). However, it was evident from this analysis that all size ranges were adequately sampled across the size range (Supplemental Figure 2) so no data were excluded. The UVP5 reports the exact depth at which a particle is recorded, however to estimate abundance, observations must be binned over fixed depth intervals. Our deployments had variable descent depths and speeds with more casts descending to 500m than 1000m and descents quicker through the epipelagic than the mesopelagic (see Barth and Stone (2022) for an extended discussion of UVP5 data processing). For the present study, Rhizaria abundances were estimated in 25m vertical bins, which offer a moderate sampling volume per bin (average 0.948𝑚^3^ in the epipelagic and 0.589𝑚^3^ in the mesopelagic; Supplemental Table 1) while still maintaining ecologically relevant widths. However, concentrations in a 25m bin would need to be greater than 2.428 ind. 𝑚^−3^and 3.912 ind. 𝑚^−3^, in the epipelagic and mesopelagic respectively, to fall below a 10% non-detection risk (Barth and Stone, 2024; Benfield et al., 1996). Because we typically observed many rhizaria taxa below these concentrations, we present the 25m binned data to visualize broad-scale average distributions. For quantifying and modelling Rhizaria abundances, we present integrated abundance estimates, with each cast. Due to the variable descent depths of the UVP, data are categorized as epipelagic (0-200m), upper mesopelagic (200-500m), and lower mesopelagic (500-1000m). The average sampling volume integrated through these regions were 7.59𝑚^2^, 7.06𝑚^2^, and 11.77𝑚^2^, with non-detection thresholds at 0.30 ind. 𝑚^−2^, 0.33 ind. 𝑚^−2^, and 0.20 ind. 𝑚^−2^ respectively. All UVP data processing was done using the EcotaxaTools package in R (Barth 2023).

### Modelling environmental controls of Rhizaria Abundance

Generalized Additive Models (GAMs) were used to assess the relationship between integrated Rhizaria abundance and different environmental factors. GAMs offer the ability to model non-linear and non-monotonic relationships, which can be particularly useful in assessing ecological relationships (Wood, 2017) and have been successfully applied to Rhizaria ecology (Biard and Ohman, 2020). The mgcv package (Wood, 2001) was used to construct models relating environmental parameters to each taxonomic group’s integrated abundance estimates from each cast. To select the most parsimonious model for each analysis, a backwards step-wise approach was taken. First, a full model was fit using any term which may be ecologically relevant.

Terms were fit using maximum likelihood with a double penalty approach on unnecessary smooths (Marra and Wood, 2011). The smoothness parameter was restricted (k = 6) to prevent overfitting the models. At each iteration of the backwards step-wise procedure, the model term with the lowest F score (least statistically significant) was removed. This was repeated until all model terms were statistically significant or 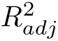 was substantially reduced. Models were fit for each region; epipelagic, upper mesopelagic, and lower mesopelagic. In cases where observations were too sparse for a given taxonomic grouping, models were not run. All code and full models are available in code, as well as intermediate data products at https://github.com/TheAlexBarth/RhizariaSeasonality.

## Results

### Environmental Variability

The BATS sampling region is southeast of Bermuda, situated in the oligotrophic North Atlantic Subtropical Gyre. Due to the sampling location, while the environmental conditions are generally low in variation and oligotrophic, there is considerable influence from deep winter-mixing and summer stratification as well as secondary influences from mesoscale eddies which result in spatiotemporal heterogeniety (Lomas et al., 2013; McGillicuddy et al., 1998). Variability in the water column structure was visible during the study period (Figure 2). This is best evidenced through the temperature profiles; In the late summer and early fall there was a stratified water column with high temperatures in the surface (<75m) (Figure 2A) and slightly elevated salinity (Figure 2B). This warm, stratified period appeared more intense during the few months sampled in 2019. In 2021, we observed the stratified layer slowly dissipated into the winter months following mixed layer entrainment. There was a consistent oxygen minimum zone (OMZ) located at about 800m deep (Figure 2C). February 2021 saw a notable downwelling event, likely due to a passing anti-cyclonic eddy which impacted the local region during early 2021. During this phaes, warmer, oxic water was significantly depressed deeper into the mesopelagic. This process was reversed in the spring months (March, April) primary due to the interaction of convective mixing and the passing of a strong cyclonic eddy resulting in a deep cold mixed layer. Primary production was highest during the spring mixing period, evidenced both by in-situ fluorescence (Figure 2D) and productivity incubation experiments (Figure 3A). Originating near the surface, the productivity peak moved deeper throughout the spring and declined into the summer (Figure 2D). However, there was a notable, yet smaller productivity bump in the late summer and early fall (Figure 3A) which occurred deeper in the epipelagic (Figure 2D). The particle concentration (184𝜇𝑚 - 450𝜇𝑚) was closely coupled to chlorophyll-a patterns.

**Figure 2:**
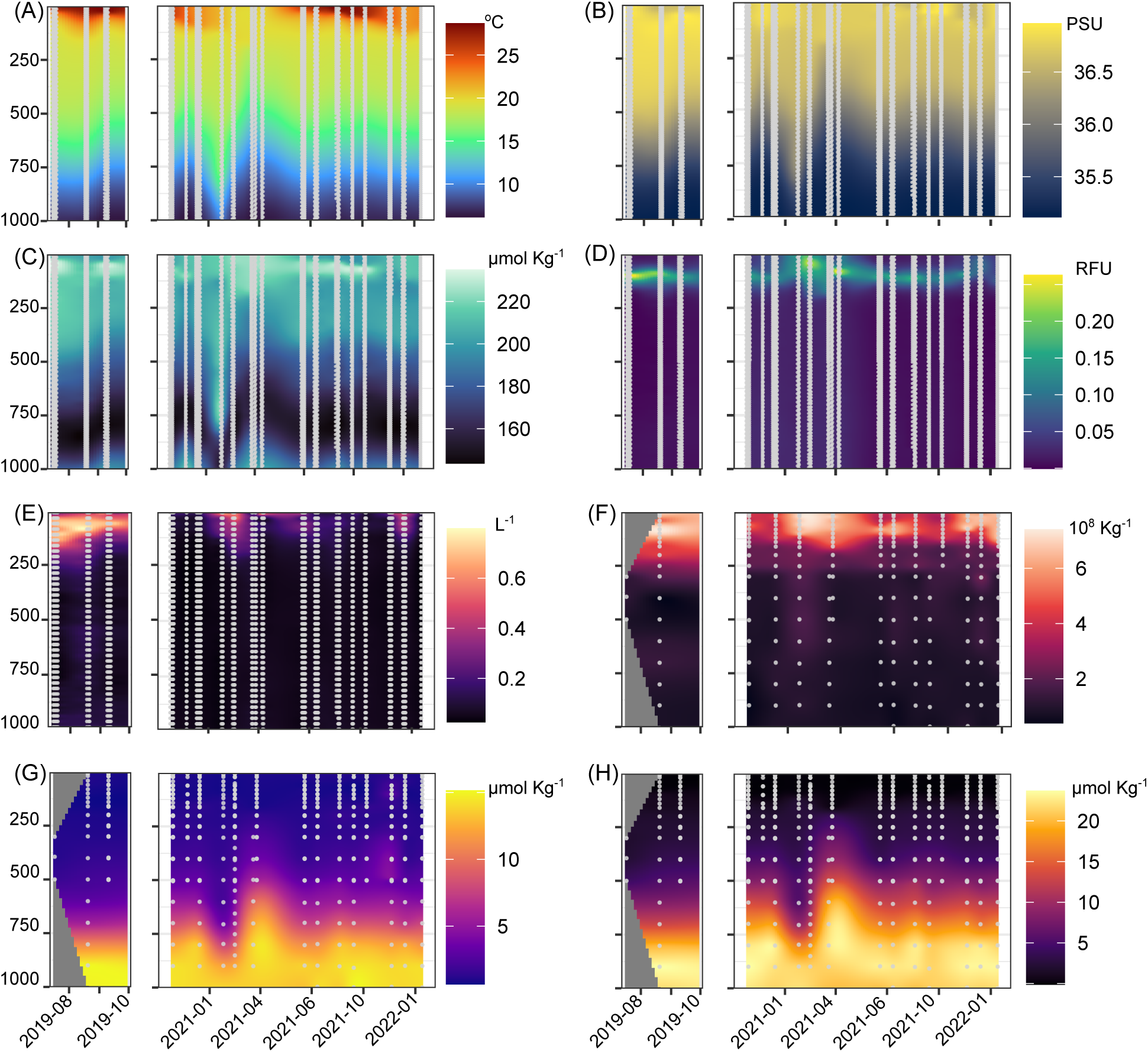
Environmental profiles across time-series of study period. Y axis shows depth in meters. (A) Temperature. (B) Salinity. (C) Dissolved Oxygen. (D) In-situ chlorophyll fluorescence. (E) Particle concenctrion (184 - 450𝜇𝑚). (F) Bacteria Abundance. (G) Silica. (H) Nitrate

**Figure 3:**
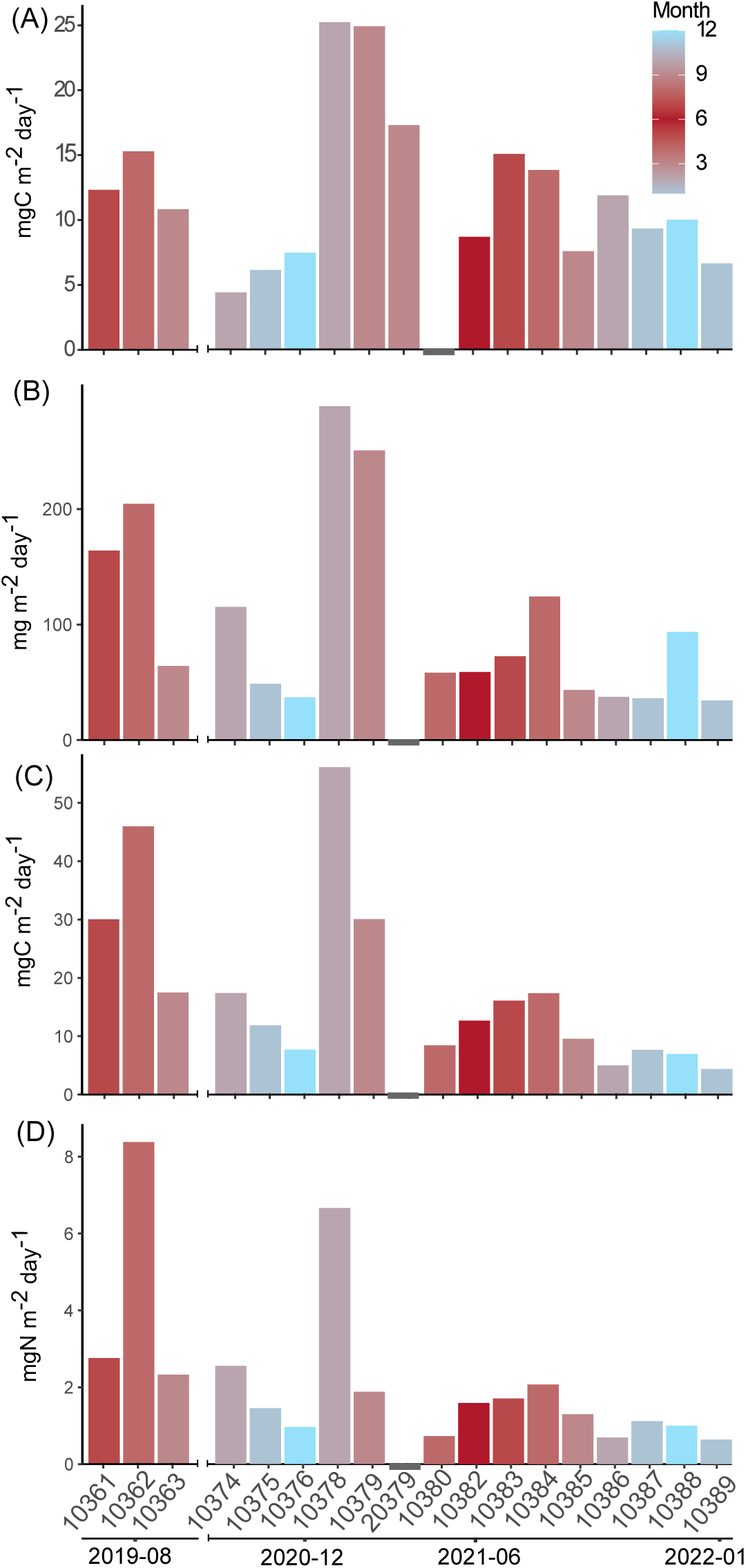
Primary productivity (A) and flux estimates from total mass (B), carbon (C), and nitrogen (D). Values are from monthly cruises with month display1e2d in corresponding colors. Absent data are shown by grey bar.

Overall, particle concentration was high near the surface during the 2021 spring bloom, then moved deeper throughout the water column attenuating throughout the lower epipelagic (Figure 2E). Similarly, heterotrophic bacteria abundance was closely linked to overall productivity, althrough there was a more consistent moderate-abundance layer near the top of the mesopelagic (∼250m) (Figure 2F). Concurrent with the secondary fall production peak, there was also higher particle concentration and bacterial abundance in the later summer and early fall. Interestingly, while primary productivity estimates from July-August were not that different between 2019 and 2021 (Figure 3A), chlorophyll-a florescence, particle concentration, and bacterial abundance were much higher in 2019’s summer/fall (Figure 2D-F). Inorganic nutrients (𝑆𝑖 and 𝑁𝑂_3_) were generally well stratified, with low concentrations in the epipelagic and increasing throughout the mesopelagic. However, both nutrients did vary vertically in accordance with the 2021 February downwelling and spring mixing period (Figure 2G-H). Additionally, in the late fall of 2021, 𝑆𝑖 concentrations were slightly elevated in the mid-mesopelagic (Figure 2G).

Overall mass flux to the mesopelagic was highest during the 2021 February downwelling (Figure 3B). Generally, export was similarly high during March, declining in April then increasing slightly throughout the summer and early fall. While magnitude was slightly different, this pattern was consistent with total mass, carbon and nitrogen fluxes (Figure 3B-D). Higher mass, carbon and nitrogen flux also occurred in the 2019 late summer - early fall period.

### Rhizaria abundance and distribution

Across all imaged mesozooplankton (>900𝜇𝑚), Rhizaria comprised a considerable fraction of the total community. Considering the total abundances of the observational period, Rhizaria comprised on average, 42.6% of all mesozooplankton abundance (Supplemental Figure 3). Copepods were the second most abundant, comprising 35.5% and all other living mesozooplankton were 22%. The large contribution of Rhizaria to the mesozooplankton community is most prominent in the epipelagic, where they accounted for 47% of all meso-zooplankton. In the mesopelagic rhizaria were a smaller (but still prominent fraction), at 38% in the upper layers (200-500m) and 37% in the deeper mesopelagic (500-1000m).

Total average Rhizaria abundance had a bimodal distribution with respect to depth. Total abundance was highest just below the surface (0-100m), with secondary, wider peak occurring in the mid mesopelagic (Figure 4A). Variation in depth binned abundance was large, likely due to seasonal variability but also increased from the detection-risk described in the methods. The vertical distribution pattern and abundance varied considerably across taxonomic groups. Radiolaria were some of the most abundant taxa observed, particularly in the epipelagic (Figure 4, Figure 5B). This pattern was led by Collodaria, whose colonies were abundant in the upper epipelagic and declined into the top of the mesopelagic (Figure 4C). Acantharea displayed a bimodal distribution accounting for a large portion of the total Rhizaria pattern (Figure 4B, Figure 5). Foraminifera had a similar bimodal distribution, yet their overall average densities were much lower and spread wider throughout the mesopelagic (Figure 4E). Phaeodaria families exhibited a wide range of vertical distribution patterns. The most abundant, Aulacanthidae, also had a bimodal pattern but the density was highest in the lower mesopelagic (Figure 4D). Aulosphaeridae had low average density and was nearly homogeneously distributed throughout the water column, although slightly lower in the epipelagic (Figure 4F). Castanellidae were the only Phaeodaria who appeared to be effectively restricted to the photic zone (Figure 4G). Alternatively, Coelodendridae primarily occurred in the lower mesopelagic (Figure 4H). A few individuals from the families Tuscaroridae and Medusettidae were also observed in the mesopelagic, yet they were much rarer (data not shown).

**Figure 4:**
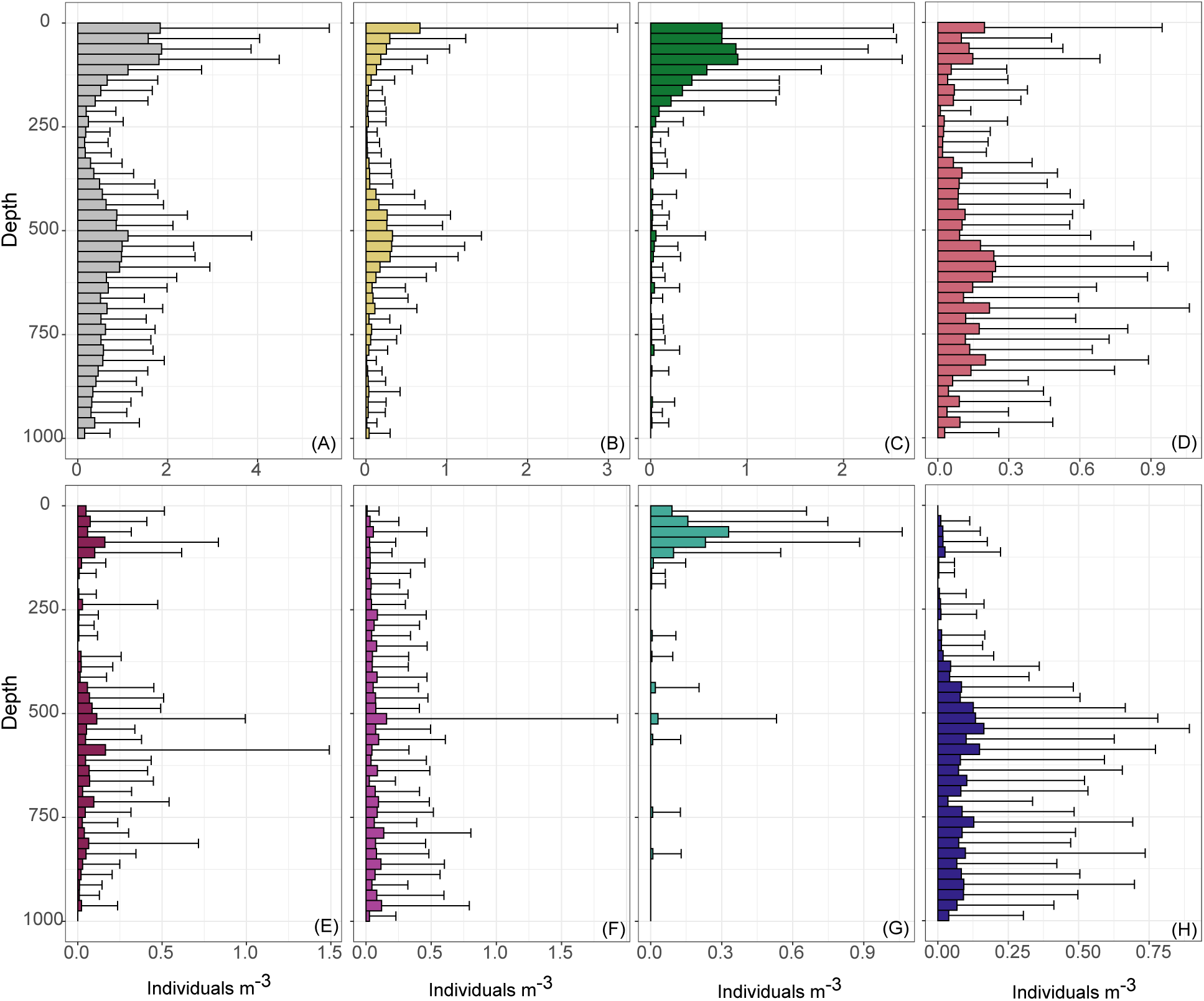
Average abundance of Rhizaria in 25m bins, across entire study period. Shown are total Rhizaria (A), Acantharea (B), Collodaria (C), Aulacanthidae (D), Foraminifera (E), Aulosphaeridae (F), Castanellidae (G), Coelodendridae (H).

**Figure 5:**
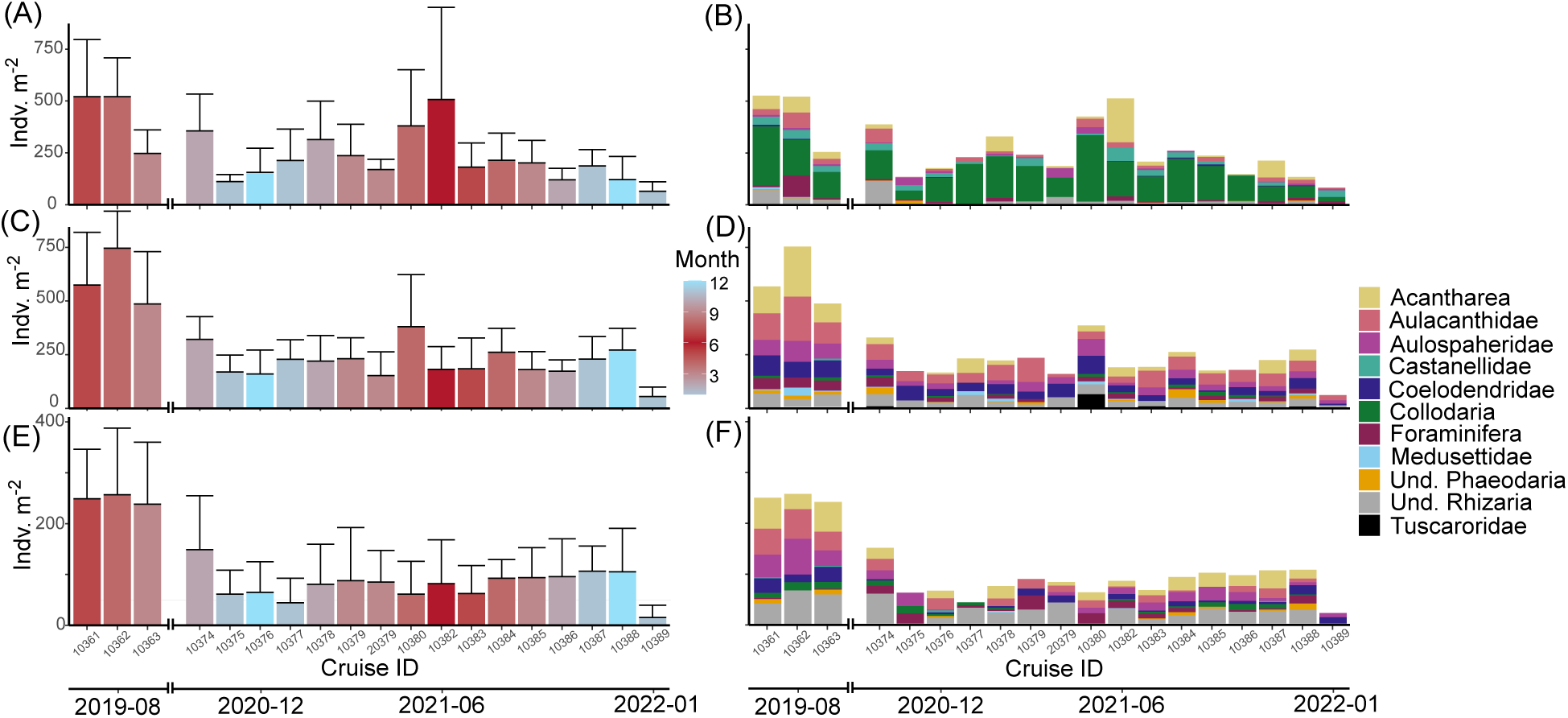
Seasonality of Rhizaria integrated abundance for the epipelagic (0-200m) (A-B), upper mesopelagic (200-500m) (C-D), lower mesopelagic (500-1000m) (E-F). Left panels (A,C,E) display total integrated abundance per monthly cruise colored by month. Right panels (B, D, F) display community composition of each total abundance.

Between the monthly cruises, Rhizaria integrated abundance varied in the epipelagic. Highest average abundance occurred in June 2021 and was lowest during the winter months (Figure 5A). The 2019 later summer - fall period also had much higher integrated abundance than similar months in 2021. While the majority of integrated abundance in the epipelagic was consistently attributable to Collodaria, Acanthrea abundance occurred sporadically and could account for a large portion of the total in some months (Figure 5B). The mesopelagic integrated abundance was much more consistent across monthly cruises, although average abundance was notably higher in 2019 (Figure 5C-F). The community composition in the mesopelagic was more diverse, mostly comprised of Phaeodarias. However, Acantharea and unidentified Rhizaria also were common members of the community (Figure 5D, 5F).

### Body size throughout the water column

Very few taxa had consistent distributions throughout the water column. Only Acanthrea, Foraminifera, Aulacanthidae, and Aulosphaeridae were consistently abundant in the epipelagic and mesopelagic. To investigate if the populations or morphologies shifted throughout the water column, we compared the sizes (Equivalent Spherical Diameters, ESD) between mesopelagic and epipelagic groups for each taxa. All groups were significantly different on average (Wilcox Rank Sum p-value <0.001). Acantharea were smaller, on average in the mesopelagic while all other taxa tended to be larger (Figure 6).

**Figure 6:**
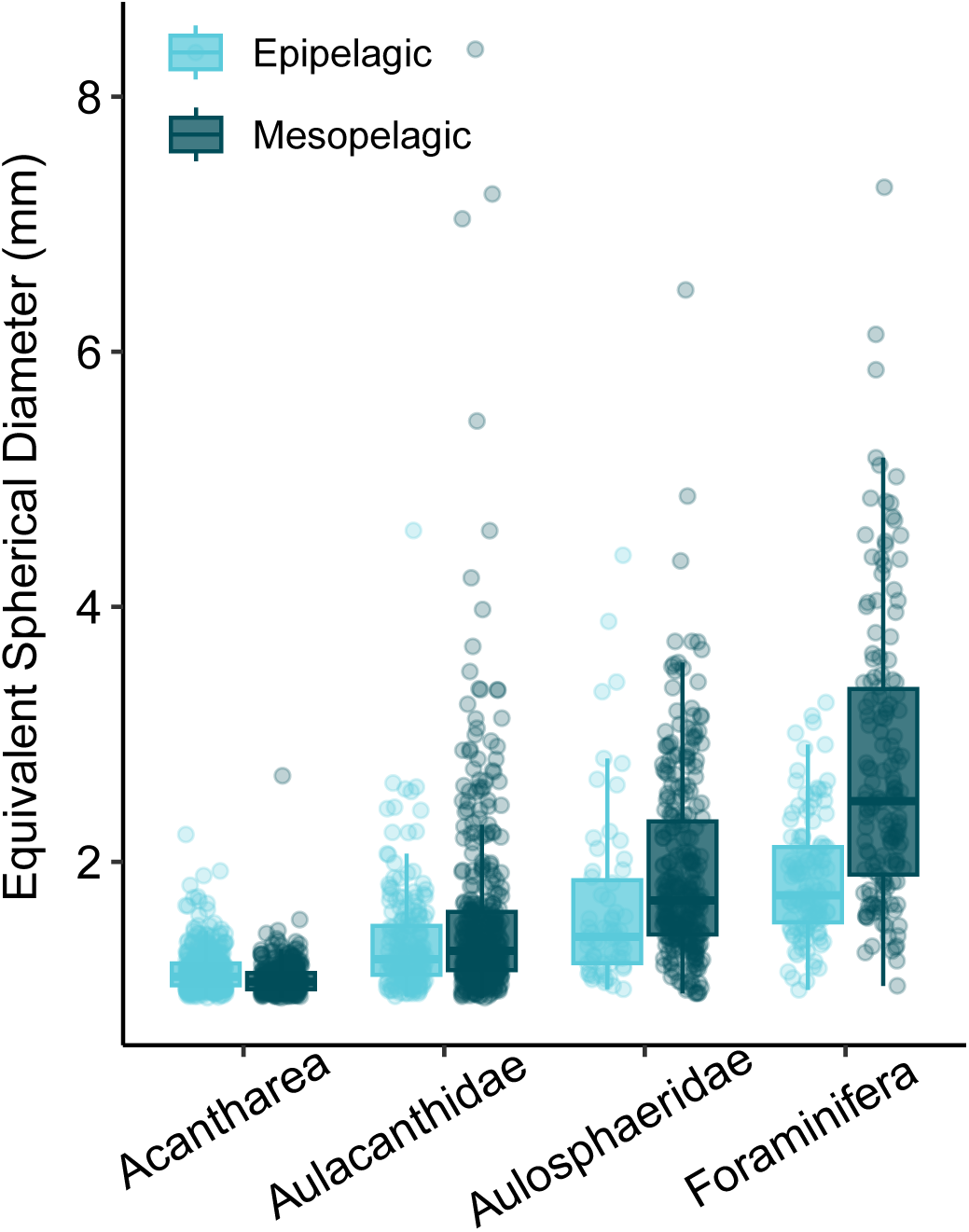
Comparison of average sizes (ESD) amongst Rhizaria taxa which occurred throughout the water column.

### Environmental Drivers of Rhizaria Abundance

For total Rhizaria integrated abundance the GAMs produced moderate fits (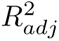= 0.406-0.603) (Table 1). In the epipelagic, there were several significant predictor variables including inorganic nutrients (𝑁𝑂_3_and 𝑆𝑖), water quality parameters (Salinity, DO), primary production, and particle concentration (Table 1). However, the upper and lower mesopelagic were exclusively explained by particle-related variables (concentration and mass flux) (Table 1).

**Table 1:**
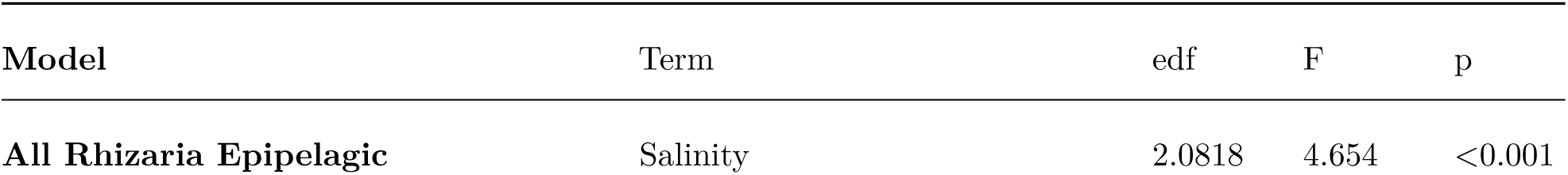
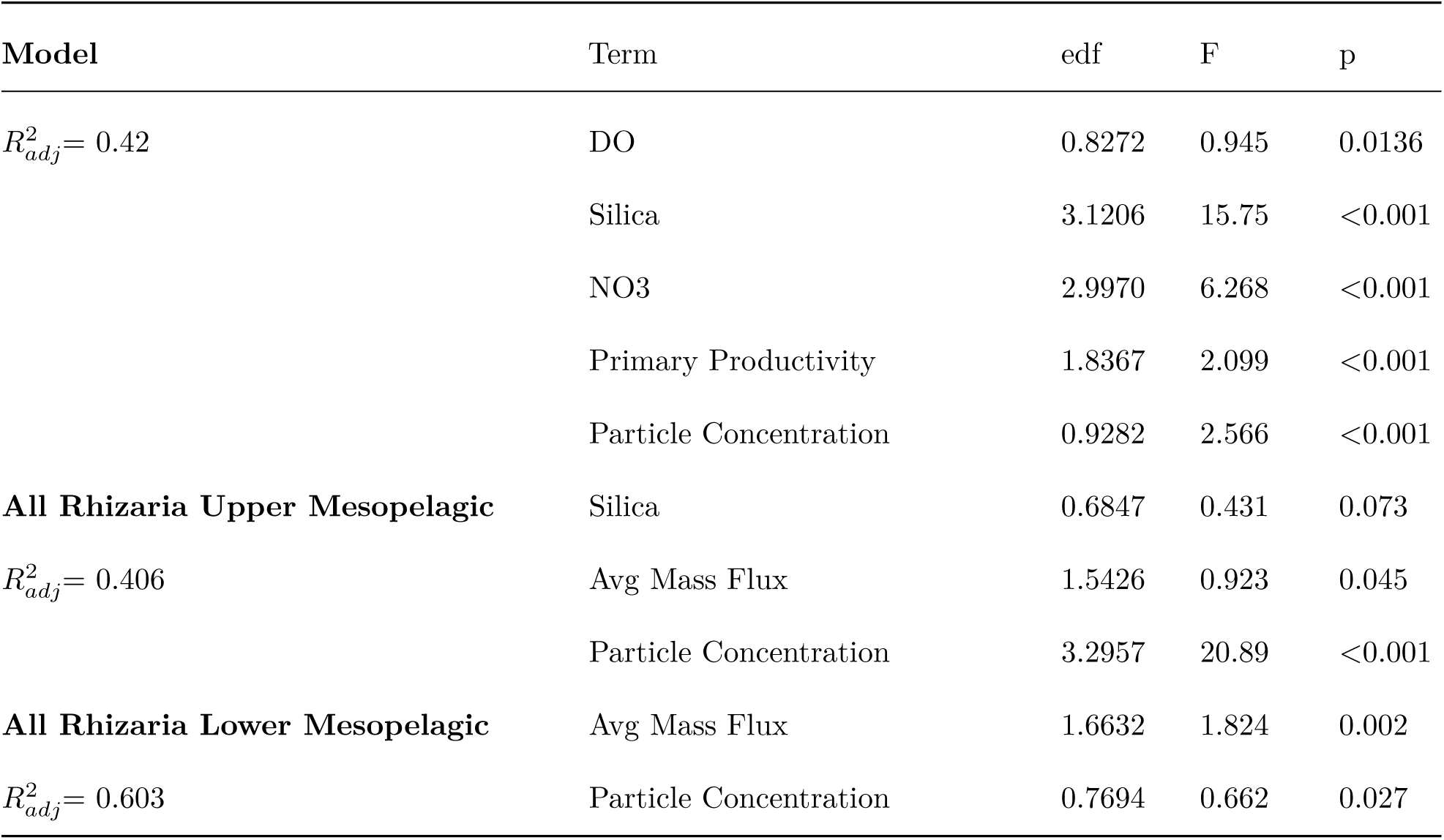
Generalized Additive Model results for integrated total Rhizaria abundance in different regions of the water column.

GAMs for individual taxa were much less consistent in their fits (Table 2). This is likely in part due to the high number of non-observations for certain taxa. Note that due to low abundances, GAMs were not constructed for Tuscaroridae or Medusettidae. Furthermore no, significant terms were found for a model with Aulosphaeridae in the epipelagic nor Foraminifera in the mesopelagic.

**Table 2:** Taxa-specific generalized additive models for different regions of the water column.

| Model | Term | edf | F | p |
| --- | --- | --- | --- | --- |
| **Acantharea Epipelagic**<br>R2adj=0.53 | Salinity | 2.264 | 4.113 | <0.001 |
|  | O2 | 2.579 | 6.712 | <0.001 |
|  | Avg Mass Flux | 4.810 | 22.81 | <0.001 |
|  | Bacteria #/L | 1.599 | 1.317 | 0.0117 |
|  | Particle Concentration | 0.853 | 1.158 | 0.0087 |
| **Acantharea Upper Mesopelagic**<br>R2adj=0.231 | Avg C Flux | 1.296 | 0.621 | 0.0439 |
|  | Avg N Flux | 1.619 | 1.283 | 0.0026 |
|  | Particle Concentration | 0.952 | 3.886 | <0.001 |
| **Acantharea Lower Mesopelagic**<br>R2adj=0.509 | Temperature | 2.076 | 4.155 | <0.001 |
|  | Avg N Flux | 1.494 | 1.216 | 0.0113 |
|  | Primary Productivity | 0.766 | 0.648 | 0.0253 |
|  | Particle Concentration | 2.037 | 10.03 | <0.001 |
| **Aulacanthidae Epipelagic**<br>R2adj=0.251 | Salinity | 0.792 | 0.757 | 0.0241 |
|  | Avg N Flux | 0.869 | 1.324 | 0.0018 |
|  | Primary Productivity | 2.312 | 7.704 | <0.001 |
|  | Particle Concentration | 2.008 | 2.200 | 0.0017 |
| **Aulacanthidae Upper Mesopelagic** | Particle Concentration | 2.832 | 9.802 | <0.001 |
| R2adj = 0.158 |  |  |  |  |
| **Aulacanthidae Lower Mesopelagic** | Avg Mass Flux | 1.622 | 1.991 | 0.002 |
| R2adj=0.298 | Particle Concentration | 2.123 | 6.706 | <0.001 |
| **Aulosphaeridae Upper Mesopelagic** | Particle Concentration | 2.653 | 13.06 | <0.001 |
| R2adj=0.2 |  |  |  |  |
| **Aulosphaeridae Lower Mesopelagic** | Particle Concentration | 0.972 | 6.248 | <0.001 |
| R2adj=.147 |  |  |  |  |
| **Castanellidae Epipelagic** | Avg C Flux | 1.462 | 0.816 | 0.0421 |
| R2adj = 0.124 | Primary Productivity | 0.771 | 0.665 | 0.0247 |
|  | Particle Concentration | 3.956 | 4.623 | <0.001 |
| **Coelodendridae Upper Mesopelagic** | Avg N Flux | 0.822 | 0.919 | 0.0183 |
| R2adj=.113 | Particle Concentration | 0.970 | 6.208 | <0.001 |
| **Coelodendridae Lower Mesopelagic** | Particle Concentration | 1.773 | 4.873 | <0.001 |
| R2adj=.133 |  |  |  |  |
| **Collodaria Epipelagic** | Salinity | 0.925 | 2.267 | <0.001 |
| R2adj=0.16 | O2 | 2.015 | 2.217 | 0.002 |
|  | Bacteria #/L | 1.843 | 2.100 | 0.002 |
| **Foraminifera Epipelagic** | Temperature | 4.414 | 8.789 | <0.001 |
| R2adj=0.445 | O2 | 1.603 | 1.287 | 0.0111 |
|  | Avg N Flux | 2.512 | 3.396 | <0.001 |
|  | Primary Productivity | 0.824 | 0.920 | 0.0153 |
|  | Particle Concentration | 0.919 | 2.257 | <0.001 |

Epipelagic Acantharea were explained by several predictor variables and had a good fit (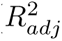 = 0.53, Table 2). Most notable smooths were mass flux and particle concentration, which had a weak positive association (Figure 7A), with July 2021 as a clear outlier where Acantharea abundances were high in the epipelagic despite lower fluxes and particle concentrations (Figure 5). Foraminifera had a good fitting GAM in the epipelagic (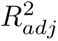 = 0.445). There were several significant explanatory variables, although the clearest pattern was observed of high temperatures associated with more Foraminifera abundance (Table 2, Figure 7B).

**Figure 7:**
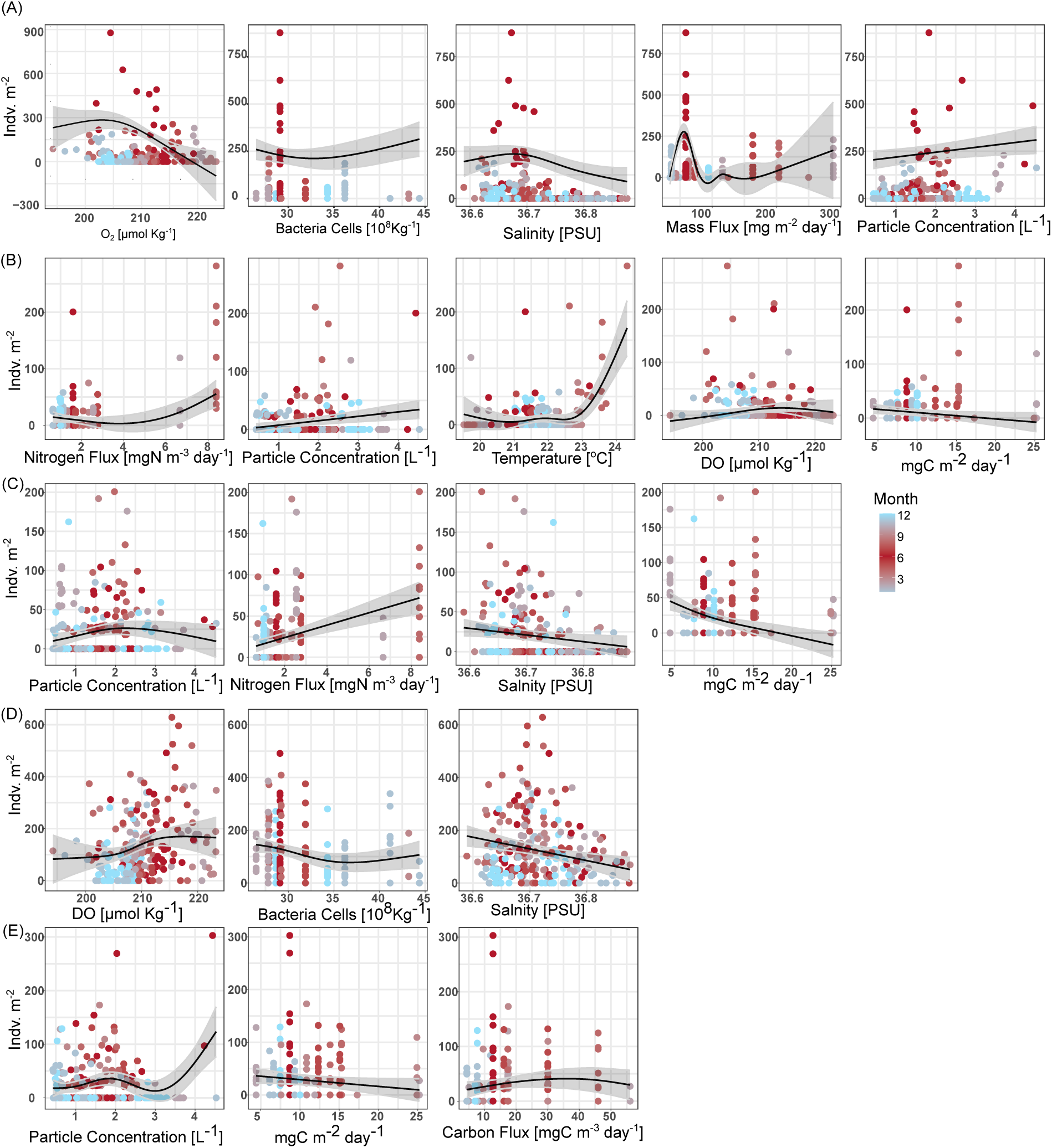
Partial effects of smooth terms in taxa-specific GAM models from the epipelagic (0-200m). Effects are grouped by taxa; Acantharea (A), Foraminifera (B), Aulacanthidae (C), Collodaria (D), Castanellidae (E).

Epipelagic Aulacanthidae similarly had several predictor variables which were significant, including both water quality parameters and particle/flux predictors (Table 2). Interestingly, Aulacanthidae had primary production as a significant predictor, yet there was not a clear association (Figure 7C). There was a fit for Collodaria in the epipelagic (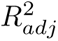 = 0.16), although there was a logit-like relationship where higher abundances tended to occur during higher DO conditions in the surface waters (Figure 7D). Castanellidae also had similarly poor fits in the epipelagic (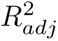 = 0.124) (Table 2, Figure 7E).

In the upper mesopelagic (200-500m), abundances were generally low (Figure 4) so GAMs were only constructed for Acantharea, Aulacanthidae, Aulosphaeridae, and Coelodendridae (Table 2). All these models had generally poor fits (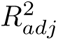 < 0.25). Yet, for all upper mesopelagic models, particle concentration was a significant explanatory variable (Table 2, Figure 8). Carbon flux was significant for Acantharea and nitrogen flux was significant for both Acantharea and Coelodendridae (Table 2, Figure 8A,D). The lower mesopelagic also had generally poor GAM fits for taxa specific models (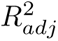 <0.3), with the exception of Acantharea (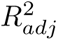 = 0.509). Acantharea in the lower mesopelagic was most clearly positively associated with particle concentration and nitrogen flux, as well as temperature to a slight degree (Figure 9A). For all Phaeodarias with a significant model, particle concentration was a main predictor variable (Table 2, Figure 9B-D). Aulacanthidae had the best fitting model of the Phaeodarias (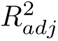 = 0.298), which also included mass flux as a statistically significant smooth (Figure 9B).

**Figure 8:**
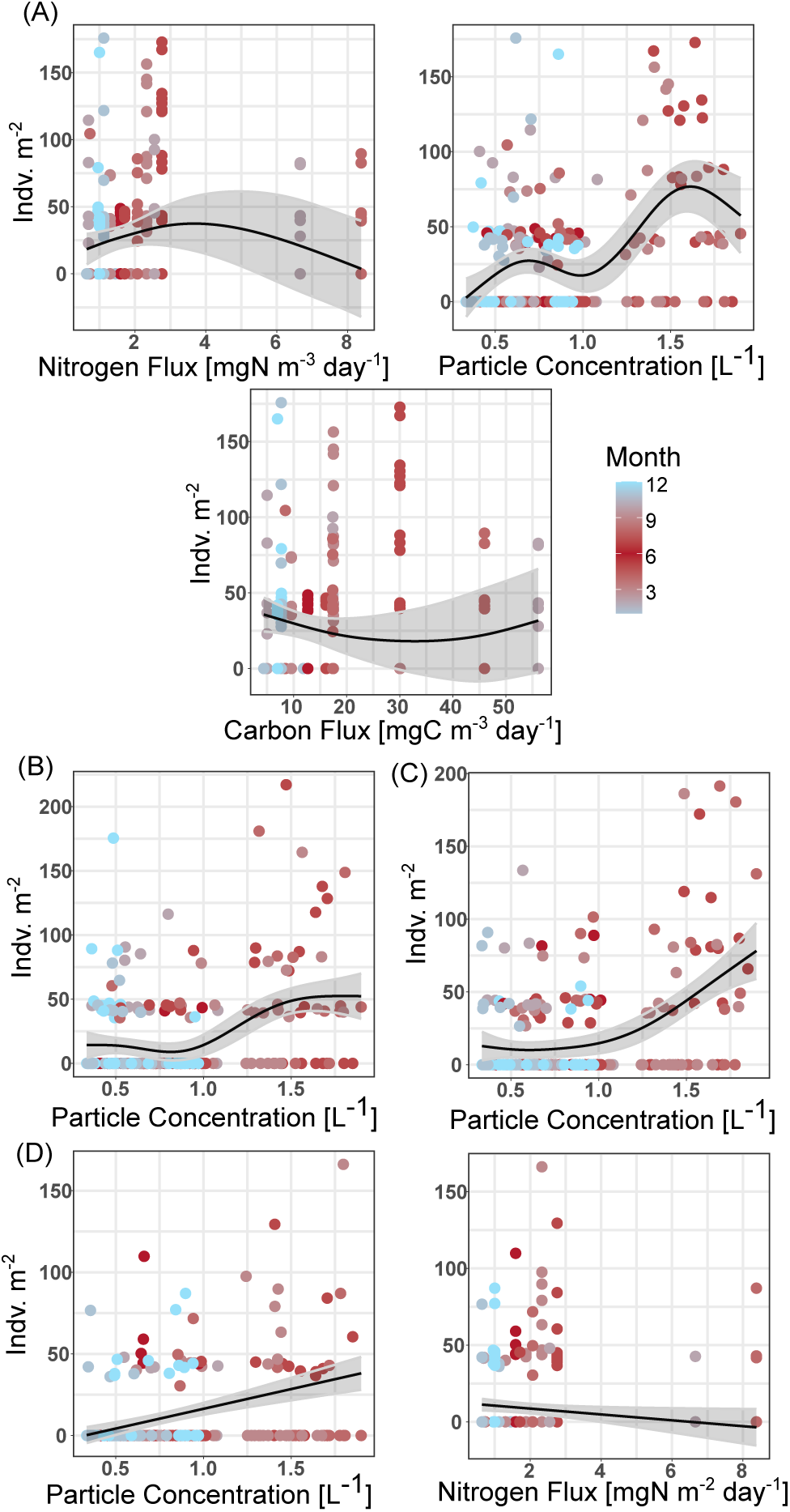
Partial effects of smooth terms in taxa-specific GAM models from the upper mesopelagic (200-500m). Effects are grouped by taxa; Acantharea (A), Aulacanthidae (B), Aulosphaeridae (C), Coelodendridae (D).

**Figure 9:**
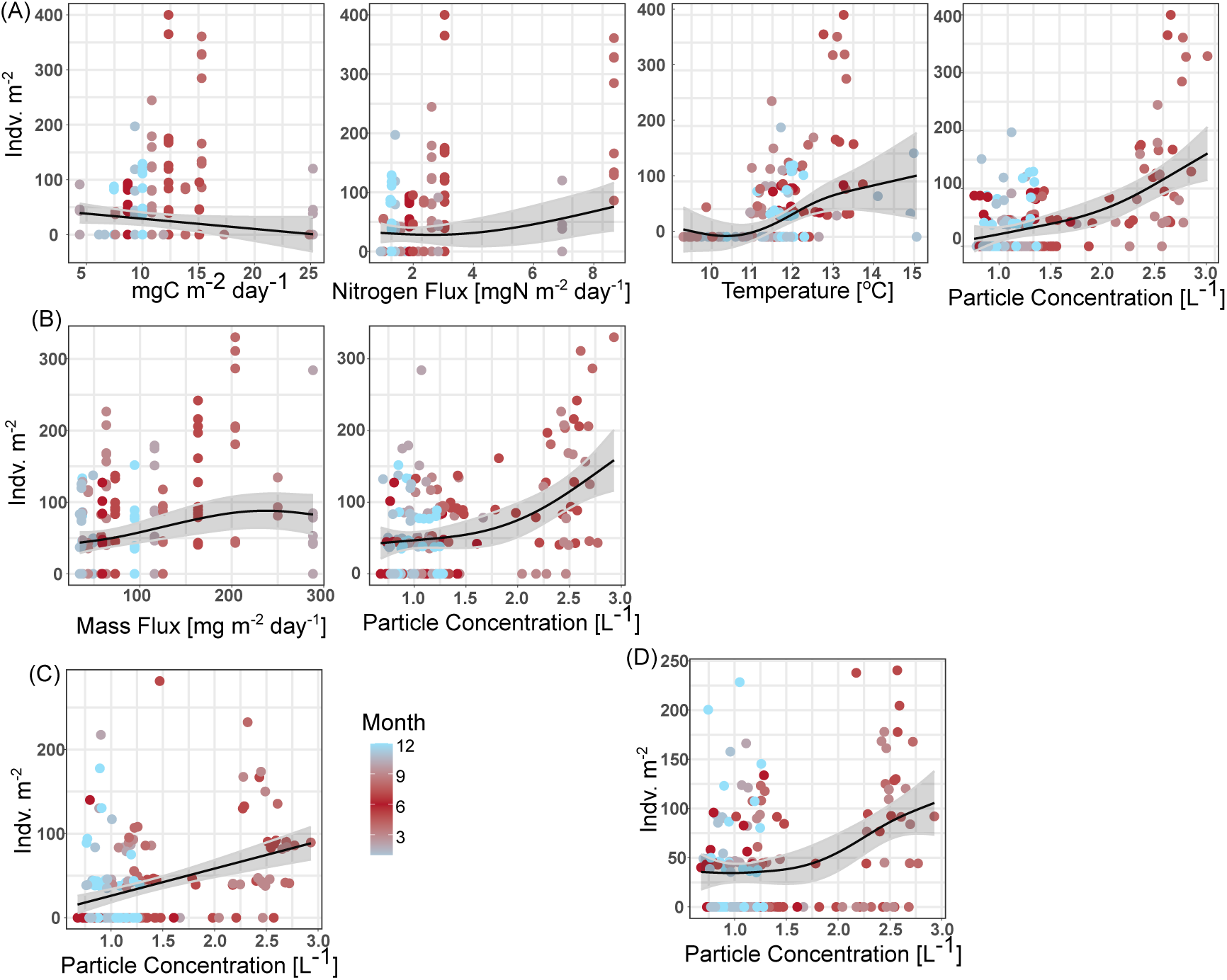
Partial effects of smooth terms in taxa-specific GAM models from the lower mesopelagic (500-1000m). Effects are grouped by taxa; Acantharea (A), Aulacanthidae (B), Aulosphaeridae (C), Coelodendridae (D).

## Discussion

### Overall Rhizaria abundance and patterns

In the epipelagic Rhizaria exhibited a notable seasonal pattern. Rhizaria abundances were higher in the summer months and lower during the winter. During a prior time period, Blanco-Bercial et al. (2022) noted that there is considerable seasonality in the community composition of all protists. Despite the seasonality of total Rhizaria abundance, community composition was relatively consistent, with Collodaria representing the bulk of the community. It should be noted that the overall taxonomic resolution of the UVP5 is fairly low, so there may be a switching of species within the broad groups identified in this study which were not captured. Throughout the mesopelagic, month-to-month variation in 2021 was relatively low. Again, this is consistent with observations from metabarcoding of the whole protist community in the same study region (Blanco-Bercial et al., 2022). This finding is not surprising as the overall seasonal variation in environmental conditions in this region were low.

Overall Rhizaria were the most commonly identified group of mesoplankton throughout the study period. We do note that the UVP5 commonly captures *Trichodesmium* colonies, yet these were excluded in this comparison as they are strictly autotrophs. It should be noted that previous work has suggested that avoidance behavior with the UVP is possible, at times likely, for visual and highly mobile zooplankton (Barth and Stone, 2022). Thus, the percent contribution reported here (42.7%) of Rhizaria to the total mesozooplankton community may be inflated due to under sampling of organisms such as Euphausiids and Chaetognaths which have quick escape responses. Regardless, it is worth noting that in the same region, with data collected in 2012 and 2013 using similar calculation methods, Biard et al. (2016) estimated Rhizaria only contribute 15% of the total mesozooplankton community in the upper 500m. Likely, Rhizaria display considerable interannual variability. In the present study, we noticed considerably higher Rhizaria abundance throughout the water column in 2019 compared to 2021. While this may have been driven by increased mass flux, more information is needed to truly understand the magnitude by which Rhizaria can vary interannually.

### Relationship to environmental parameters

In general, the fit of most GAMs were moderate to poor. One possible reason for the poor fits may have been that for some taxa, conditions were not variable enough to capture a range of conditions at which they may exist. For instance, Collodaria were the most abundant taxa observed, yet the fit of their GAM was particularly poor. In studies which covered a wider range of parameters, Collodaria has been shown to strongly vary with changes in parameters such as temperature, chlorophyll-a, mixing, and water clarity (Biard et al., 2017; Biard and Ohman, 2020). Alternatively, Acantharea had relatively good fitting GAMs. These taxa also had some of the largest variation from month to month on cruises. Thus, it may be that in the oligotrophic, the relatively stable conditions can support certain taxa while others are more sporadic. It should also be noted that due to the challenge of adequately sampling enough volume to overcome low-detection issues, GAMs were run on integrated data. However, variation with environmental parameters throughout the water column are likely, just not captured in the modelling aspect of this study. One consistent parameter which had significant positive associations was particle concentration. This observation is not surprising as most rhizaria likely to some extent engage in flux feeding.

### Vertical Structure and Trophic Roles

In this study we present a clear pattern of vertical zonation between different Rhizaria groups. Largely, the taxonomic composition and vertical positioning were similar to Rhizaria zonation in the California Current Ecosystem (Biard and Ohman, 2020). It should be noted however, that the secondary abundance peak reported in the present study is lower. This is likely due to the more oligotrophic nature of the study region, were the euphotic zone penetrates deeper into the water column. Most prevalent in the epipelagic were Collodaria. These mixotrophic Radiolaria have long been reported to contribute to primary productivity in the euphotic zone (Dennett et al., 2002; Michaels et al., 1995). Collodaria are thought to be particularly successful globally in oligotrophic regions due to their photosymbiotic relationships (Biard et al., 2017, 2016). We observed the highest abundance of Collodaria during June 2021, supporting the notion they can thrive during the typically low-nutrient conditions of summer stratification. However, Collodaria also increased during the spring mixing period, suggesting that they can thrive during conditions which may typically be thought to favor autotrophs. Furthermore, while Collodaria were primarily absent from below 250m, there were a few instances of deeper colonies being observed. Global investigations of polycystine flux, suggest that deep-Collodaria in Oligotrophic regions may be a consequence of isothermal submersion (Boltovskoy, 2017). Alternatively, surface waters at BATS often mix into the mode water during the seasonal mixing, so Collodaria in the deeper waters may be a result of diapyncal mixing. Another effectively exclusively epipelagic Rhizaria was the Phaeodaria family of Castanellidae. All Phaeodaria are thought to be fully heterotrophic (Nakamura and Suzuki, 2015), nonetheless a number of studies, including this one, report Castanellidae to be typically found in the lower epipelagic (Biard et al., 2018; Biard and Ohman, 2020; Zasko and Rusanov, 2005). It should be considered that perhaps Castanellidae specializes in feeding on sinking particles directly at the base of the epipelagic. Given its smaller size (Nakamura and Suzuki, 2015), Castanellidae does not need a large diameter to efficiently flux feed at the typically particle rich region of the lower epipelagic. Both Castanelldiae and Collodaria had poor fitting GAMs. This is somewhat of a surprise for Collodaria who had large abundance. However, given the consistency of their abundance, it may be that this study did not capture a wide enough range of conditions for describing Collodaria’s preferred niche.

The mesopelagic generally was home to known heterotrophic organisms, particularly for those which were constrained to exclusively occupy deeper waters. This is consistent with Blanco-Bercial et al. (2022)’s observation of an auto-/mixotroph to heterotroph gradient in the protist community. The upper mesopelagic interestingly had relatively low total abundance. This low-abundance region likely reflects the dynamics of productivity and export throughout the water column. While productivity and thus sinking particles for flux feeders are high in the euphotic zone, much of this is attenuated throughout the epipelagic. So, while the base of the epipelagic may provide a rich feeding environment for Castanellidae, smaller protists, or heterotrophic bacteria (Figure 2F), the region from 200-500m might be otherwise food poor. Perhaps it is more advantageous for Rhizaria to situate deeper, in darker regions of the twilight zone. Also it should be noted that Phaeodaria utilize silica to build their opaline tests, and silica concentrations started to increase around 500m (Figure 2G). Although 𝑆𝑖 was not a significant smooth for any taxa-specific model, this lack of association might be due to the overall lack of variation of 𝑆𝑖 between integrated abundance of each cast. Aulosphaeridae was only found to have significant relationships, although weak fits, to particle concentration in the mesopelagic. In our study, while consistently observed, overall abundances of Aulosphaeridae were very low. In the Pacific Ocean, on California’s Coast, much higher abundances of Aulosphaeridae have been reported (Biard and Ohman, 2020; Zasko and Rusanov, 2005) and they have massive potential to impact silica export (Biard et al., 2018). Coelodendridae were also seemingly restricted to the deeper section of the mesopelagic. This is interesting given that in the California Current, (Biard and Ohman, 2020) found a bimodal distribution in Coelodendridae. There are several morphotypes corresponding to different taxa of Coelodendridae (Biard and Ohman, 2020; Nakamura and Suzuki, 2015). So it may be that only a few types of Coelodendridae were observed in this study, while the epipelagic variety was not. Alternatively, the lower epipelagic of the California Current may provide adequate habitat for Coelodendridae, which is not available in the oligotrophic Sargasso Sea.

A number of taxa were found to have a bimodal distribution, with considerable populations in both the epipelagic and mesopelagic. Aulacanthidae had a bimodal distribution, although abundances were highest in the lower mesopelagic. Foraminifera also had a bimodal distribution. Some lineages of Foraminifera are known to host photosymbionts (Biard, 2022a; Kimoto, 2015), however they are also efficient predators commonly seen throughout the mesopelagic (Caron and Be, 1984; Gaskell et al., 2019). Thus it is not surprising to find their presence in both locations of the water column. Foraminifera are also known to vary their vertical distribution across their life cycle in phase with lunar cycles (Biard, 2022a; Bijma et al., 1990; Gaskell et al., 2019; Kimoto, 2015). However, the sampling scheme of the BATS program does not capture this frequency and was not investigated in the present study.

Acantharea also had a bimodal distribution, with much larger abundances than Aulacanthidae or Foraminifera. Most prior studies of Acantharea vertical distribution found them concentrated in near surface layers of the water column Zasko and Rusanov (2005). This would support the paradigm that large Acantharea abundances may be supported by their mixotrophic abilities (Michaels et al., 1995; Suzuki and Not, 2015). While the UVP5 images cannot distinguish between mixotrophic and heterotrophic Acantharea, the GAMs constructed for Acantharea abundance found positive associations with particle concentration and mass flux, suggesting a higher reliance on heterotrophy. Recently Mars Brisbin et al. (2020) described apparent predator behavior amongst near-surface Acantharea. Thus it is likely that epipelagic Acantharea may commonly be heterotrophic. Yet, it should be noted in the Sargasso Sea, both heterotrophic and symbiotic lineages of Acantharea have been reported (Blanco-Bercial et al., 2022). Additionally, Michaels (1988) noted that the majority of Acantharea (by abundance) were smaller than 160𝜇𝑚. While that estimate may be inflated due to inability to capture larger cells, small Acantharea were not captured in the present study. Thus, trophic strategy may shift based on sizes of Acantharea.

Decelle et al. (2013) proposed a hypothetical life cycle for cyst-bearing (strictly heterotrophic) Acantharea. This hypothesized life cycle suggests that epipelagic Acantharea are adult populations, which form cysts that sink into the mesopelagic, then reproduce and rise. Furthermore, given that horizontal transfer of symbionts between generations of Acantharea is unlikely due to their spawning behavior, the newly spawned mesopelagic Acantharea are not necessarily required to rapidly return to the photic zone (Decelle et al., 2013, 2012). This hypothesis predicts that Acantharea in the mesopelagic would be smaller (Decelle et al., 2013). Mars Brisbin et al. (2020) provided some support for this hypothesis, with a significant decrease in Acantharea sizes with depth. Although the authors also observed low abundances in the mesopelagic and noted that the smaller sizes may be due to lower food availability (Mars Brisbin et al., 2020). Since food is more scarce in the mesopelagi, nutritional quality lower, yet flux feeders would likely grow larger to increase their feeding range (Biard and Ohman, 2020). In the data collecting in this study, Acantharea in the mesopelagic were significantly smaller than the epipelagic, despite the other bimodal taxa (Foraminifera and Aulacanthidae) being significantly larger with depth. This provides added support for the hypothesis that cyst-forming Acantharea may utilize different sections of the water column throughout their life cycle. However to further investigate this, more work is needed with higher temporal and taxonomic resolution.

### Conclusions and Considerations

This study provides a detailed look at Rhizaria abundance over time throughout the water column in a major oligotrophic gyre. We show that their abundances are generally related to particle concentration and flux, although lack of environmental variability may have reduced the fit of our GAMs. Considering the potential role of Rhizaria in the biological carbon pump, they may have a somewhat mixed role. In the shallower regions, where smaller Rhizaria are abundant, they may be an attenuating force on sinking particles (Stukel et al., 2019). It should be noted that in our study, we focused on the “small” particle concentration field (<450𝜇𝑚), and these particles are generally slower sinking than large particles. However, once consumed and repackaged by larger Rhizaria, they can sink quicker and contribute more to overall flux (Michaels, 1988). Thus, Rhizaria may act as an aggregation mechanism. However, this is largely speculation, to truly test this, more work is needed measuring Rhizaria flux.

The vertical partitioning documented in this study do support the hypothesis that mixotrophic rhizaria will occupy shallower waters while deeper waters are dominated by heterotrophy. However the degree to which mixotrophic Rhizaria in the euphotic zone rely on heterotrophy versus symbiosis is uncertain. Collodaria were recorded as consistent and dominant members of the near surface region. These organisms have the potential to contribute considerably to the otherwise low productivity of oligotrophic regions. However, their role in food webs is not well understood. While this study represents a step-forward in our understanding of Rhizaria ecology, continued research on Rhizaria is much needed to better understand their ecology. Particularly extended descriptive work to capture interannual patterns. Also work defining biotic interactions, feeding rates, productivity, and life history are all rich fields of interest in Rhizaria.

