## Supplemental for "Rhizaria in the oligotrophic ocean exhibit clear temporal and vertical variability"

Supplemental Table 1. UVP sampling volume summary statistics. Epi indicates epipelagic (0-200m) while meso indicates mesopelagic (200-1000m).

| **Cruise ID** | **Year** | **Month** | **Zone** | **Casts** | **Mean Vol per 25m bin (m**^-3^) | **Integrated Mean Volume (m**^-2^) | **Total Volume (m**^-3^) |
| --- | --- | --- | --- | --- | --- | --- | --- |
| 10361 | 2019 | Jul | epi | 19 | 1.043 | 8.348 | 154.436 |
| 10361 | 2019 | Jul | meso | 19 | 0.598 | 7.170 | 329.223 |
| 10362 | 2019 | Aug | epi | 13 | 0.943 | 7.547 | 94.344 |
| 10362 | 2019 | Aug | meso | 12 | 0.578 | 6.941 | 173.521 |
| 10363 | 2019 | Sep | epi | 20 | 0.981 | 7.851 | 152.116 |
| 10363 | 2019 | Sep | meso | 20 | 0.612 | 7.347 | 324.506 |
| 10374 | 2020 | Oct | epi | 17 | 0.980 | 7.838 | 133.240 |
| 10374 | 2020 | Oct | meso | 17 | 0.551 | 6.611 | 240.754 |
| 10375 | 2020 | Nov | epi | 6 | 0.909 | 7.271 | 43.628 |
| 10375 | 2020 | Nov | meso | 6 | 0.557 | 6.686 | 106.971 |
| 10376 | 2020 | Dec | epi | 22 | 0.958 | 7.663 | 168.583 |
| 10376 | 2020 | Dec | meso | 22 | 0.581 | 6.975 | 365.012 |
| 10377 | 2021 | Jan | epi | 8 | 0.705 | 5.637 | 45.098 |
| 10377 | 2021 | Jan | meso | 7 | 0.520 | 6.238 | 86.812 |
| 10378 | 2021 | Feb | epi | 12 | 0.967 | 7.736 | 92.827 |
| 10378 | 2021 | Feb | meso | 12 | 0.591 | 7.088 | 204.357 |
| 10379 | 2021 | Mar | epi | 6 | 0.921 | 7.365 | 44.189 |
| 10379 | 2021 | Mar | meso | 6 | 0.570 | 6.835 | 87.715 |
| 20379 | 2021 | Mar | epi | 6 | 0.891 | 7.127 | 39.200 |
| 20379 | 2021 | Mar | meso | 6 | 0.549 | 6.584 | 74.074 |
| 10380 | 2021 | Apr | epi | 7 | 0.663 | 5.301 | 33.130 |
| 10380 | 2021 | Apr | meso | 6 | 0.553 | 6.630 | 99.457 |
| 10382 | 2021 | Jun | epi | 29 | 0.949 | 7.592 | 220.173 |
| 10382 | 2021 | Jun | meso | 29 | 0.559 | 6.706 | 422.495 |
| 10383 | 2021 | Jul | epi | 20 | 1.037 | 8.298 | 165.968 |
| 10383 | 2021 | Jul | meso | 20 | 0.561 | 6.738 | 284.660 |
| 10384 | 2021 | Aug | epi | 14 | 0.895 | 7.160 | 95.764 |
| 10384 | 2021 | Aug | meso | 14 | 0.600 | 7.199 | 209.978 |
| 10385 | 2021 | Sep | epi | 10 | 1.002 | 8.013 | 78.130 |
| 10385 | 2021 | Sep | meso | 10 | 0.670 | 8.045 | 174.312 |
| 10386 | 2021 | Oct | epi | 13 | 0.971 | 7.765 | 93.182 |
| 10386 | 2021 | Oct | meso | 13 | 0.594 | 7.127 | 179.358 |
| 10387 | 2021 | Nov | epi | 18 | 0.919 | 7.351 | 132.321 |
| 10387 | 2021 | Nov | meso | 18 | 0.635 | 7.616 | 259.564 |
| 10388 | 2021 | Dec | epi | 21 | 0.951 | 7.607 | 159.745 |
| 10388 | 2021 | Dec | meso | 21 | 0.631 | 7.569 | 363.945 |
| 10389 | 2022 | Jan | epi | 9 | 0.924 | 7.389 | 64.650 |
| 10389 | 2022 | Jan | meso | 9 | 0.611 | 7.332 | 129.537 |

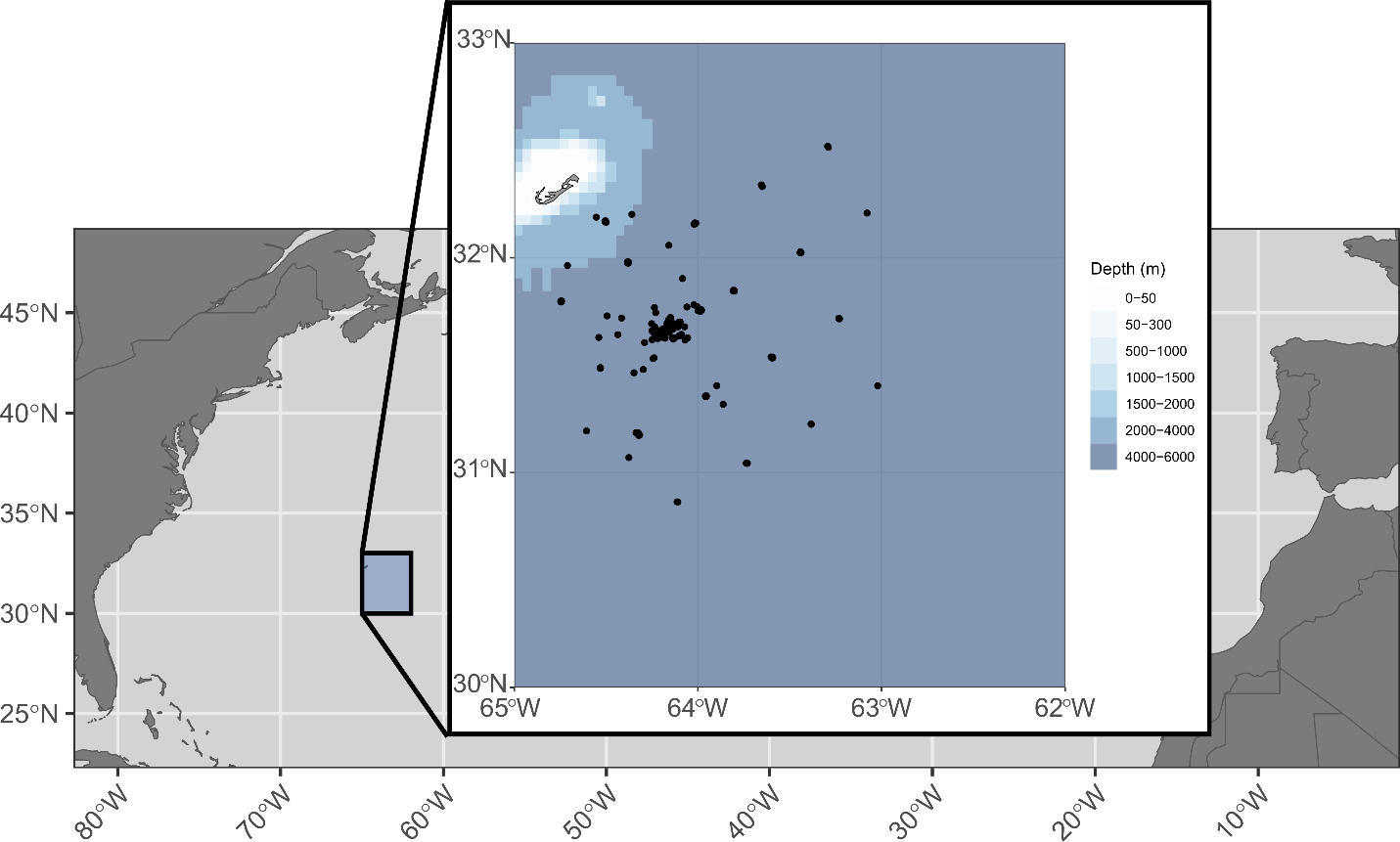

Supplemental Figure 1. Map of casts during study period. Bathymetry data from ggOceanMaps R package: Vihtakari M (2024). ggOceanMaps: Plot Data on Oceanographic Maps using 'ggplot2'. R package version 2.2.0, https://mikkovihtakari.github.io/ggOceanMaps/.

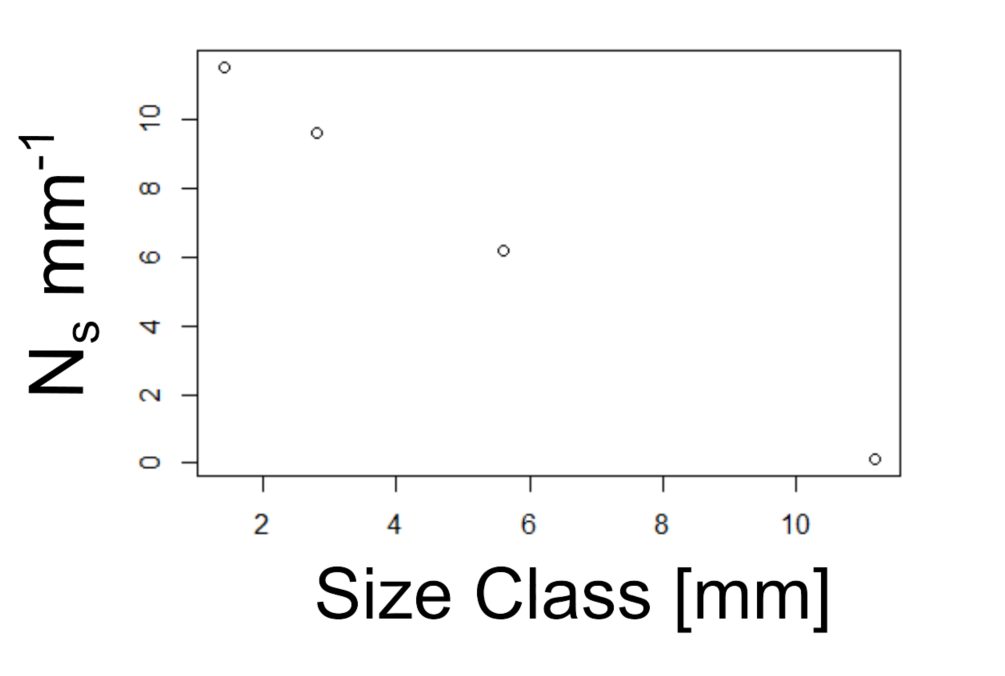

Supplemental Figure 2. Numerical size spectra of all Rhizaria vignettes.

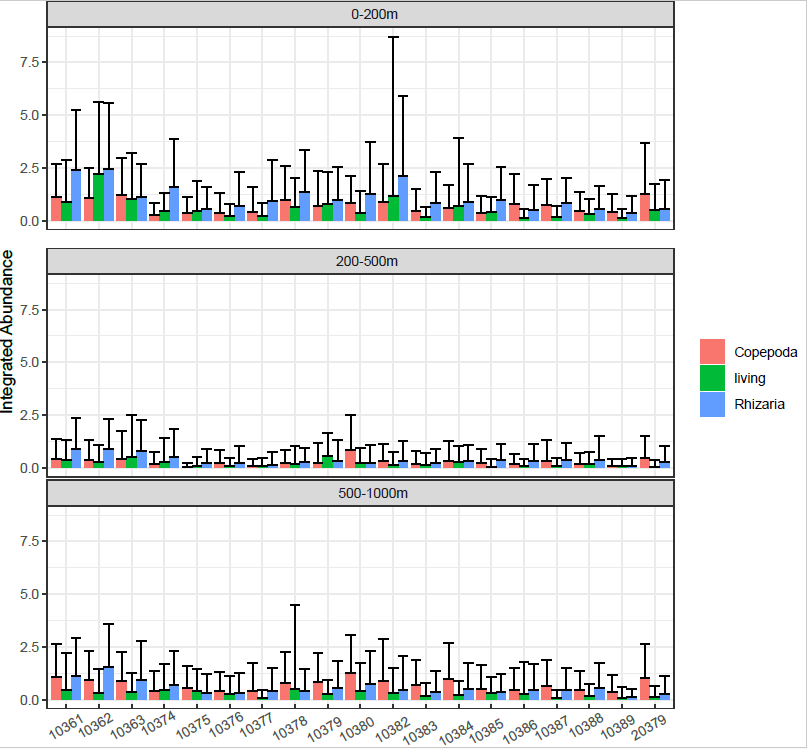

Supplemental Figure 3. Integrated abundances across cruises for different oceanic regions; epipelagic (0-200m), Upper mesopelagic (200-500m), lower mesopelagic (500-1000m). Shown are different color bars corresponding to taxonomic grouping. Living indicates all other living mesozooplankton.
